# Malaria-induced remodelling of the bone marrow microenvironment mediates loss of haematopoietic stem cell function

**DOI:** 10.1101/477190

**Authors:** Myriam L.R. Haltalli, Samuel Watcham, Nicola K. Wilson, Kira Eilers, Alexander Lipien, Maria L. Vainieri, Nicola Ruivo, Heather Ang, Robert E. Sinden, Ken R. Duffy, Berthold Göttgens, Andrew M. Blagborough, Cristina Lo Celso

## Abstract

Severe infections are a major source of stress on haematopoiesis, where consequences for haematopoietic stem cells (HSCs) have only recently started to emerge. HSC function critically depends on the integrity of complex bone marrow niches, which have been shown to be altered during ageing and haematopoietic malignancies. Whether the bone marrow (BM) microenvironment plays a role in mediating the effects of infection on HSCs remains an open question. Here we used an murine model of malaria coupled with intravital microscopy, single cell RNA-Seq, mathematical modelling and transplantation assays to obtain a quantitative understanding of the proliferation dynamics of haematopoietic stem and progenitor cells (HSPCs) during *Plasmodium* infection. We uncovered that during *Plasmodium* infection the HSC compartment turns over significantly faster than in steady state, and that a global interferon response and loss of functional HSCs are linked to alterations in BM endothelium function and osteoblasts number. Interventions that targeted osteoblasts uncoupled HSC proliferation and function, thus opening up new avenues for therapeutic interventions that may improve the health of survivors of severe infections.

## INTRODUCTION

Haematopoietic stem cells (HSCs) maintain the turnover of all blood cells throughout our lifetime. They reside within the bone marrow (BM), where they interact with several stroma and haematopoietic cell types. These complex niches regulate HSC function by maintaining their quiescence and by supporting their self-renewal when proliferation occasionally occurs (Morrison & Scadden, 2014). Once out of the niche, HSC progeny differentiate into highly proliferative haematopoietic progenitor cells (HPCs), which further develop into erythroid, megakaryocyte and immune myeloid and lymphoid lineages.

Infection is a natural stressor of the haematopoietic system that results in the consumption of immune cells that require replacement. In parallel with the development of innate and adaptive immune responses, infections and inflammatory cytokines stimulate haematopoietic stem and progenitor cells (HSPCs) to modify the composition of their progeny to cope promptly with the increased demand for mature cells (Essers *et al.*, 2009; Esplin *et al.*, 2011; King & Goodell, 2011; MacNamara *et al.*, 2011). A range of dramatic HSPC responses have been noted as a result of these insults, including an expansion of the early progenitor compartment (identified as Lineage^-^ c-Kit^+^ Sca-1^+^, or LKS), and alterations in cycling properties, long-term function and migration patterns of HSCs (Baldridge *et al.*, 2010; Rashidi *et al.*, 2014; Matatall *et al.*, 2016; Vainieri *et al.*, 2016). These phenomena all take place within the bone marrow (BM) microenvironment, yet very little is currently known about the response of HSC niche cells to infection and inflammation, and what their role may be in regulating the observed changes in haematopoietic dynamics.

Malaria is a severe and life-threatening disease that continues to cause significant morbidity and mortality, particularly across Africa, Asia and South America. In 2016, there were an estimated 216 million new cases of malaria worldwide, with approximately 445,000 deaths (WHO, 2017). *Plasmodium* infects and destroys red blood cells, and multiple studies have investigated its impact on erythropoiesis (Dörmer *et al.*, 1983; Maggio-Price, Brookoff & Weiss, 1985; Wickramasinghe *et al.*, 1989; Boehm *et al.*, 2018). It is generally accepted that malaria survivors have compromised haematopoietic and immune function in the short- and long-term (Orf & Cunnington, 2015; White, 2018), and recent studies have begun to link this to alteration in cells of myeloid lineages (Orf & Cunnington, 2015; Mamedov *et al.*, 2018). It is likely that these effects can be traced back to changes in the earliest stages of haematopoiesis, including the HSC compartment. However, so far little published work directly addresses the effect of *Plasmodium* infection on more primitive HSPCs. To fully assess the role of malaria in the progression of these pathologies and to potentially devise therapeutic strategies that preserve HSC function, further understanding of the impact of malaria on HSCs is necessary. In particular, the HSC niche has recently emerged as a valuable therapeutic target in the context of leukaemia-driven loss of HSC function (Hawkins *et al.*, 2016; Duarte *et al.*, 2018). We hypothesized that this may also be the case in the context of infection and therefore we sought to investigate the effects of *Plasmodium* infection on the HSC niche and whether they may mediate some of the effects on HSCs.

Using a murine experimental model based on the natural route of sporozoite-mediated infection, with mice receiving bites from *Plasmodium berghei*-infected mosquitoes, we previously demonstrated that multiple components of the haematopoietic tree simultaneously respond to *P. berghei* infection. We uncovered a dramatic increase in the proportion of proliferating HSCs and early progenitors at the advanced stages of blood infection (Vainieri *et al.*, 2016). This work raised questions about what cell-intrinsic and -extrinsic mechanisms mediate the haematopoietic response to *P. berghei* infection and in particular: 1) what proliferation dynamics sustain stressed haematopoiesis; 2) what molecular mechanisms drive them; 3) their effect on the long-term function of HSCs; 4) how the BM microenvironment is affected by infection; and 5) whether microenvironmental changes might contribute to the HSC phenotypes observed.

Here, we combined phenotypic, functional, molecular and intravital microscopy (IVM) analyses to gain insights into the haematopoietic and BM microenvironment responses over the course of infection, investigating causal links between the two. These data established that the entire HSPC compartment is affected and activated by *P. berghei* infection, with significant changes in HSPC proliferation rate, loss of the transcriptional HSC signature, and loss of HSC long-term function. We demonstrate that, as well as HSPCs, interferon (IFN) dramatically affects the BM microenvironment, with diffuse disruption of BM vascular integrity and loss of osteoblasts. Moreover, we identify that targeting the osteolineage with parathyroid hormone (PTH) reduces osteoblast loss, lowers local and systemic IFN levels and prevents HSC proliferation in response to *P. berghei* infection.

## RESULTS

### *P. berghei* infection affects the dynamics of the entire HSC compartment

To understand the consequences of *Plasmodium* infection on HSPCs, mice were infected by bites of mosquitos carrying the infective sporozoite stage of the murine parasite *P. berghei* (using naïve, uninfected mosquito bites as controls, in parallel) and their parasitemia monitored over time. Visible symptoms of cerebral malaria developed consistently within 8 days (Figure 1A) and therefore we focused our studies on the first 7 days post-sporozoite infection (psi). During this period, merozoites were first generated in the liver and were undetectable in peripheral blood up to day 3. Red blood cell invasion was subsequently initiated and measurable parasitaemia increased over day 5 and 7 psi (Figure 1A). To monitor the HSPC response to infection, we used flow cytometry analysis of the BM of infected and control mice. Striking changes developed over time and were most prominent at day 7 psi (Figure 1B), with a notable upregulation of Sca-1 expression within undifferentiated (Lineage^-^) haematopoietic cells. Consistent with our previous observations, the ratio of myeloid committed progenitors (LK cells, for Lineage^-^ c-Kit^+^ Sca-1^-^) and more primitive cells (LKS, Lineage^-^ c-Kit^+^ Sca-1^+^) was inverted, with the LK population decreasing while LKS increased. Within the LKS population, we used SLAM markers (Kiel *et al.*, 2005; Bryder, Rossi & Weissman, 2006; Foudi *et al.*, 2008; Wilson *et al.*, 2008; Grassinger *et al.*, 2010; Oguro, Ding & Morrison, 2013) to separate the cell population most enriched for HSCs from their immediate progeny – mixed lymphoid and myeloid primed multipotent progenitors (MPPs). We included Sca-1 in our gating strategies, even though its expression was upregulated, because it is widely adopted in studies focussing on the effects of infection on early HSPCs (Matatall *et al.*, 2016; Zhang *et al.*, 2016), and in our hands it was largely redundant when combined with CD150 and CD48. While the LKS CD150^+^ CD48^-/low^ phenotype enriches for HSCs, it known not to result in a pure HSC population (Wilson *et al.*, 2008; Oguro, Ding & Morrison, 2013; Cabezas-Wallscheid *et al.*, 2014; Akinduro *et al.*, 2018). Here we label that phenotype as ‘HSCs’ for brevity, but it should be considered as primitive HSPCs. By day 7 psi the ratio of primitive HSPCs (labelled HSC) to MPPs was clearly skewed towards MPPs, with a dramatic expansion of the latter population (Figure 1B).

**Figure 1:**
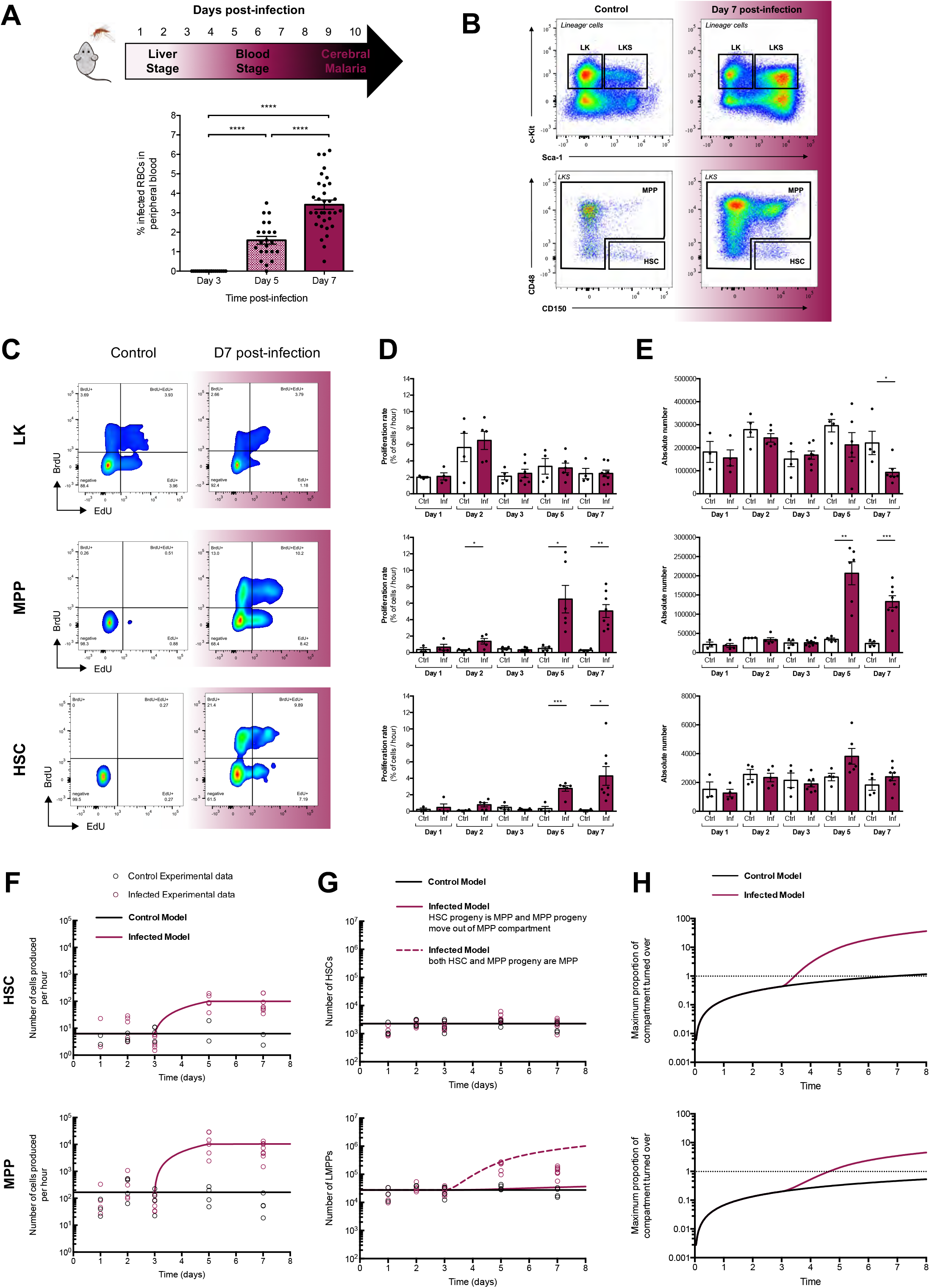
*P. berghei* infection model and analysis of the effects on HSPC populations in the BM. **(A)** Mice are exposed to control or *P. berghei* infected *A. stephensi* mosquitos at day 0. The duration of the liver and blood stages of disease, as well as the onset of cerebral complications are indicated. Parasitemia in peripheral blood over time post-infection is calculated from Giemsa-stained blood films taken from infected mice and quantified. Each dot represents one mouse. **(B)** Representative FACS plots showing the changing pattern of Sca-1 and c-Kit expression in Lineage^-^ BM haematopoietic cells. Boxes indicate the gating strategy used for LKS, myeloid-committed progenitors (LK), multipotent progenitors (MPP) and haematopoietic stem cells (HSC) compartments. **(C)** Representative FACS plots showing the gating strategy for EdU^+^ and BrdU^+^ LK, MPP and HSC populations in control and infected mice at day 7 post-infection. **(D)** Proliferation rates (proportion of the population cycling per hour) of LK, MPP and HSC populations analysed at day 1, 2, 3, 5 and 7 post-infection in control and infected mice. **(E)** Absolute numbers of LK, MPP and HSC populations analysed at day 1, 2, 3, 5 and 7 post-infection in control and infected mice. **(F)** Simple mathematical models of the number of HSCs (top panel) and MPPs (lower panel) entering S phase per hour. In the control model, the number is constant corresponding to the average of measurements from all control mice plus the infected mice up to day 3. In the infected model, the number is the same as the control model until day 3, and then linearly increases in time to the average of the data from day 5 and day 7 infected mice. Experimental data is overlaid on the graphs. **(G)** Simple mathematical models of the HSC (top panel) and MPP (lower panel) compartment sizes. For both the control and infected mice, the HSC population is modelled as a constant corresponding to the average of data from all control and infected mice. For the MPP population, the control model is a constant corresponding to the average of measurements from all control mice plus the infected mice up to day 3. In one infected model, the number is that same as the control model until day 3, and then increases corresponding to the additional number of HSCs entering S-phase per hour in the infected proliferation model beyond those in the control model. In the second infected model, the number is constant to day 3, and then increases based on the sum of the additional number of HSCs and MPPs entering S-phase per hour in the infected model beyond those in the control model. Experimental data is overlaid on the graphs. **(H)** Mathematical modelling of the maximum proportion of the HSC compartment (top panel) and MPP compartments (lower panel) turned over as a function time. Log Scale used. Dotted line visually represents the first time point at which the entire compartment could have completed one turnover. All data pooled from 3 independent infections. In **(A, D and E)** Error bars represent the mean ± s.e.m and each point represents one mouse.

To gain a quantitative understanding of the HSPC response as infection progresses, we monitored the size of multiple compartments and, as a measure of proliferation, determined the number of cells entering S-phase per hour using a dual-pulse labelling method that we recently employed to evaluate cell population dynamics in leukaemia-burdened BM (Akinduro *et al.*, 2018). Briefly, mice were first administered the thymidine analogue 5-ethynyl-2’-deoxyuridine (EdU) and, after two hours, another thymidine analogue 5-bromo-9-deoxyuridine (BrdU). Thirty minutes later, BM was harvested and analysed by flow cytometry, and cells that were labelled with BrdU alone were identified as having entered S-phase during the two-hour gap. By measuring the total number of cells within a compartment as well as the number of EdU^-^ BrdU^+^ cells, we could infer the proportion of the compartment entering S-phase per hour, i.e. the proliferation rate of each cell population analysed (Figure 1C). The resulting data revealed that the proliferation rate of LK cells remains constant throughout the course of infection. In contrast, the proliferation rates of both MPPs and HSCs were only statistically consistent with controls until day 3, before dramatically increasing by day 5 (Figure 1D). As we previously reported (Vainieri *et al.*, 2016), LK cell numbers were significantly reduced by day 7, while MPP numbers were substantially increased, and the number of HSCs remained essentially unchanged throughout the course of infection (Figure 1E). The observation that the HSC compartment does not grow in size during infection, even though the proportion of HSCs entering S-phase does, suggests that HSCs produce an increased number of differentiated progeny.

In order to investigate whether the augmented number of MPPs observed could derive solely from additional input from the HSC compartment, we performed quantitative inference via a simple mathematical model. Consistent with the flow cytometry data, in the model we assumed that both HSCs and MPPs from infected mice produce cells at the same rate as their respective controls until day 3 post-infection, followed by a linear increase in the number of cells entering S-phase per hour between days 3 and 5 (Figure 1F). This subsequently plateaus and remains constant through to day 7. In the model, the HSC compartment size is consistent with controls and constant through infection (Figure 1G, top panel), while the growth in the MPP compartment size is inferred under different assumptions, as follows. In Figure 1G, lower panel, the black line represents the control data where the MPP population size remains constant. The solid maroon line models the situation where additional HSC progeny (beyond those produced at steady-state) differentiate to become MPPs, but all MPP progeny beyond the control base-line continue to exit the MPP compartment. This model demonstrates that the HSC contribution to the MPP compartment alone is not sufficient to account for the growth of the MPP population that was observed at the later timepoints of *P. berghei* infection. The dashed maroon line (Figure 1G, lower panel) is an inference of how many MPPs should have been detected if all additional progeny resulting from the increased proliferation of *both* HSCs and MPPs gave rise to MPPs. From this analysis, it is clear that by day 7 the number of MPPs detected is lower than the model predicts. Consequently, these data are consistent with a hypothesis where the majority of the additional progeny resulting from the increase in proliferation of HSCs and MPPs up to day 5 are indeed MPPs, but after day 5 a significant proportion of the additionally produced cells must have exited the MPP compartment.

Because proliferation affects HSC long-term function, and, in particular, a small number of cell divisions is sufficient to exhaust dormant HSCs (Bernitz *et al.*, 2016), we next assessed how quickly the increase in proliferation of HSPCs could result in the whole LKS compartment being activated by *P. berghei* infection. With information on both the proliferation rate and population size of the HSC and MPP compartments, we determined the minimum time it would take for each whole compartment to turnover, under both control and infected conditions, by assuming that all cells within a compartment cycled sequentially. For control HSCs, based on the model previously described (Figure 1G), in control mice the maximum number of HSCs that could have passed through S-phase at least once within 8 days would constitute less than the whole compartment (Figure 1H, top panel, black line). However, for infected mice, by day 5 the whole compartment could have potentially already turned over once. By day 8, it is possible that the whole HSC population will have turned over approximately 5 times as a result of the faster proliferation rates beyond day 3 of infection (Figure 1H, top panel, maroon line). When similar considerations are applied to the MPP population, we calculated that this entire compartment could potentially turnover once every 8 days in control mice, but, due to the increase of entry into S-phase, this would increase to up to 40 times during the progression of infection (Figure 1H, bottom panel). These data demonstrate that *P. berghei* infection drastically impacts HSPC turnover, prompting investigation into whether phenotypic HSCs, which were still detectable in infected BM, remain functional.

### Primitive HSPCs are significantly reduced in infected mice

To test this hypothesis, we carried out transplantation assays of purified LKS SLAM primitive HSPCs from control and infected donor mice (Figure 2A-C). We injected 200 cells per recipient and followed their engraftment over 20 weeks. Cells from infected animals had dramatically reduced engraftment ability (Figure 2B). However, their multilineage potential was consistent with that of transplanted control HSCs. Upon secondary transplantation, after pooling the BM of primary recipients and purifying HSCs from control and infected groups, we observed similar levels of reconstitution ability and multilineage potential, despite a trend towards skewed myeloid lineages, was not significantly different between infected and control groups (Figure 2C). These data suggested that even though *P. berghei* infection drastically affects the overall HSC compartment and leads to a significant reduction in HSC function, a few robust and functional HSCs do remain.

**Figure 2:**
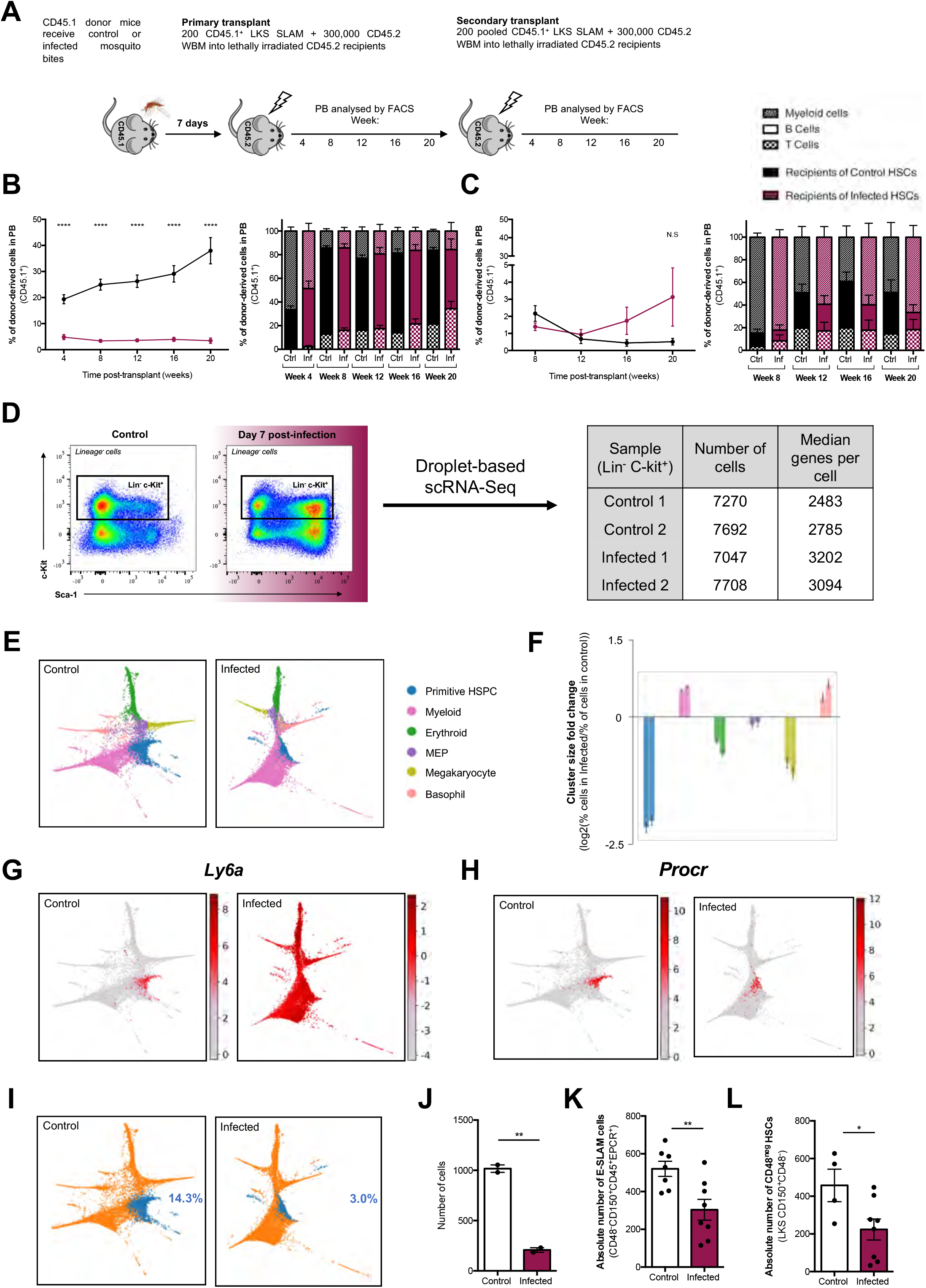
HSCs exposed to *P. berghei* undergo significant transcriptional changes as well as a loss of long-term function. **(A)** Scheme detailing the primary and secondary transplantation assay carried out to measure long-term potential of control and infected FACS purified HSCs. **(B)** and **(C)** Peripheral blood reconstitution and multilineage potential of transplanted HSCs assessed by flow cytometry post-primary **(B)** and post-secondary **(C)** transplant. Data representative of 2 independent infections and transplantation assays. N = 12 mice per group. **(D)** Sorting gate used to isolate the Lineage^-^ c-Kit^+^ cells for droplet-based scRNAseq. Table indicates the number of cells which passed QC and median number of genes detected in scRNAseq samples. **(E)** Cluster identity of control and infected cells. Control cells were grouped into 6 clusters using Louvain clustering on the k-nearest-neighbour graph. Infected cells were mapped to their corresponding control cluster and coloured accordingly, specified in the key. **(F)** Log_2_ fold-change of the percentage of control cells in each cluster divided by the percentage of infected cells mapped to that same cluster. Left-hand and right-hand bars indicate fold changes for samples from two separate mice. **(G)** Expression of *Ly6a* gene plotted on the force-directed graph embedding. **(H)** Expression of *Procr* gene plotted on the force-directed graph embedding. **(I)** (Left) Binary clustering of the control data with Primitive HSPC cells in blue and all other clusters in orange. (Right) Corresponding infected cells mapped to the control binary clusters. **(J)** Quantification of primitive HSPC cells in both control and infected samples analysed. **(K)** Absolute number of E-SLAM cells (CD48^-^CD150^+^CD45^+^EPCR^+^) in control and infected mice at day 7 post-infection, analysed by flow cytometry. Data pooled from 2 independent infections. **(L)** Absolute number of CD48^neg^ HSCs (LKS CD150^+^CD48^neg^) in control and infected mice at day 7 post-infection, analysed by flow cytometry. Data pooled from 3 independent infections. In **(B), (C), (J), (K) and (L)** Error bars represent mean ± s.e.m.

To obtain a broad and unbiased understanding of the changes taking place in HSPCs as a consequence of *P. berghei* infection, we performed single cell RNA sequencing (scRNAseq) on Lineage^-^ c-Kit^+^ cells from control and infected mice (Figure 2D). This overcame the limitations inherent to the use of the few phenotypic markers allowed by flow cytometry analyses, and allowed us to address two important questions: first, which HSPC populations remained, i.e. to what extent early haematopoiesis was reprogrammed by infection, and, second, what molecular mechanisms could be driving the observed changes. Similar numbers of cells were successfully analysed from two control and two infected animals, with 7270 and 7692 cells in control and 7047 and 7708 cells in infected samples passing quality control and a median of over 2400 genes detected per cell (Figure 2D). To understand global changes in early haematopoiesis, we grouped the Lineage^-^ c-Kit^+^ cells into 6 clusters representing different lineages and subsequently mapped the cells from infected animals against their corresponding control cluster. The data were displayed using force-directed graphs, highlighting the various lineage branches in different colours (Figure 2E). This indicated that, in infected mice, early haematopoiesis was substantially rewired towards myeloid and basophil lineages, at the expense of erythroid and megakaryocyte lineages. Quantification of the changes in proportion of specific cell types between the control and infected samples (Figure 2F) confirmed these initial observations.

Next, using binary clustering, we separated cells based on the co-expression of genes associated with the HSC state (Wilson *et al.*, 2015), including *Ly6a* and *Procr* (Figure 2G-H). As expected, *Ly6a* was overexpressed in the majority of cells sequenced from the infected mice (Figure 2G). However, the number of cells expressing a gene signature typical of the most primitive HSPCs was strikingly decreased compared to cells from control mice (Figure 2I-J). Prompted by these data, we performed flow cytometry analysis of BM cells from infected and control mice, this time focusing on immunophenotypes known to highly enrich for functional HSCs. When we combined the marker for endothelial protein C receptor (EPCR) with SLAM markers (Kent *et al.*, 2009), we detected a significant decrease in the absolute number of E-SLAM HSCs in infected mice (Figure 2K). When we focused on the LKS CD150^+^ CD48^neg^ HSC subpopulation, which was recently identified to have the highest reconstituting potential when compared to LKS CD150^+^ CD48^low^ HSCs (Akinduro *et al.*, 2018), we also observed a significant decrease in their absolute number in infected mice (Figure 2L). Together, the transplantation, scRNAseq and flow cytometry data suggest that despite the numbers of LKS SLAM cells remaining unchanged throughout infection, the function of this cell population is reduced by *P. berghei* infection. This is likely because of a reduction in the number of HSC subpopulations known to be most functional

### *P. berghei*-exposed HSPC and BM stroma cells exhibit a strong interferon response

To identify the molecular mechanisms driving the phenotypes observed, we re-examined the scRNAseq data set, asking what genes were differentially expressed between control and infected samples for each cluster. We focussed on genes that were at least 4-fold upregulated in all clusters as they were likely to drive the observed haematopoietic reprogramming. This filtering identified 20 initial genes, most of which were related to interferon (IFN), and some specifically to IFN-gamma (IFN-γ) signalling (Supplementary Figure 1 and Supplementary Table 1). We further analysed the data searching for genes whose expression correlated highly with those in the initial driving gene set (see methods), extending the list of driving genes to 109 (Supplementary Table 1). The importance of these genes was demonstrated by combining the control and infected cells into a single SPRING plot generated by including or excluding them from the analysis. When these genes were included, the control and infected groups were forced away from each other, but when they were excluded, the two groups of cells appeared much closer (Figure 3A). Interestingly, the majority of these genes were also related to IFN signalling, indicating that this is likely to be the main driver of the observed effect, yet the fact that some separation remained between the control and infected datasets suggested that even after filtering, there are still genes left that show global differential expression.

**Figure 3:**
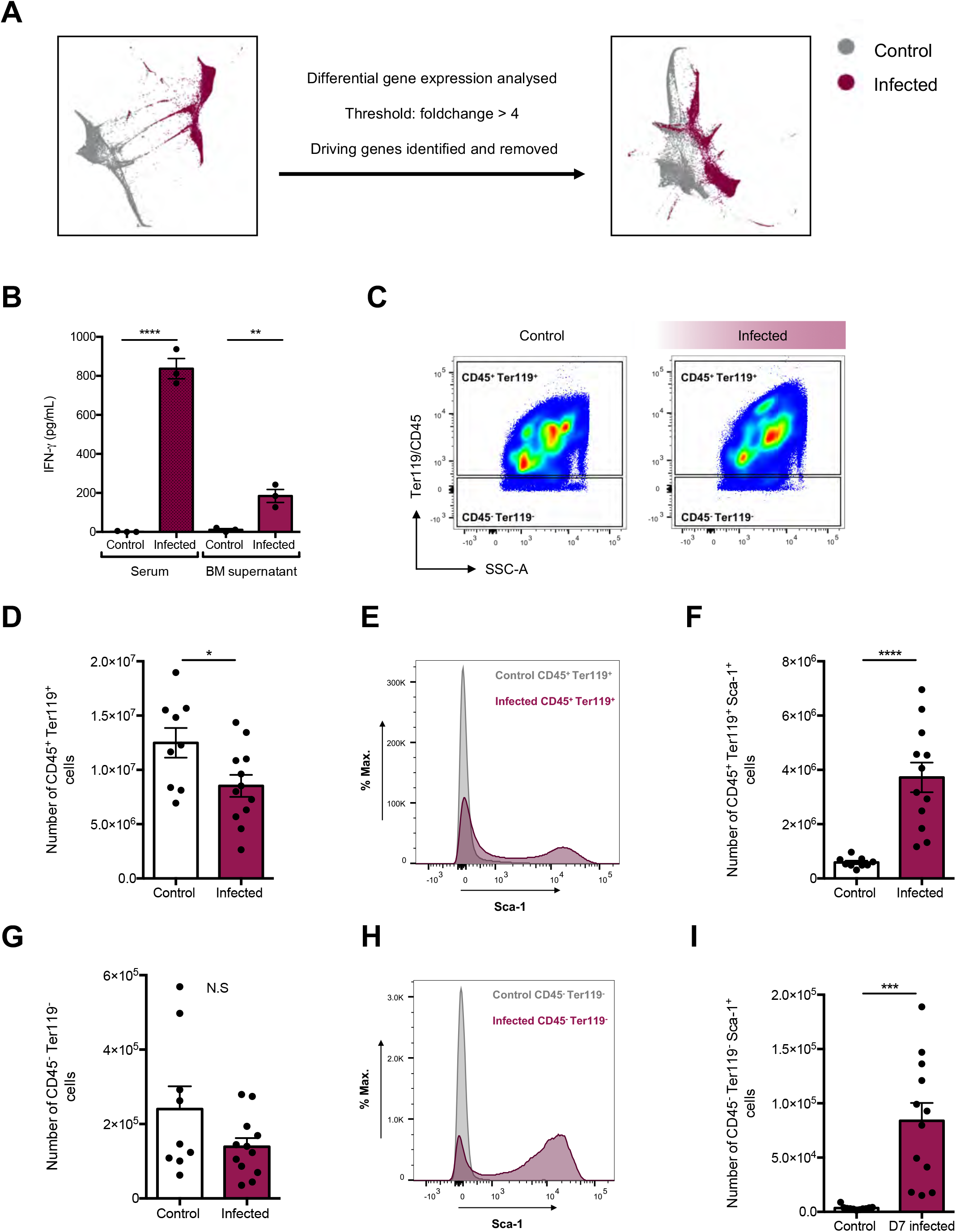
scRNAseq and analysis of the BM microenvironment reveal a significant interferon signature with *P. berghei* infection. **(A)** Plotting the combined control and infected cells with a single force-graph embedding highlights the large separation between samples according to their infection status, regardless of cell type. Recalculating the force graph with the 109 interferon genes removed suggests that these genes are responsible for the majority of the changes between control and infected cells. **(B)** IFN-γ levels in BM supernatant and serum of control and infected mice measured by ELISA. N = 3 mice per group. **(C)** Gating strategy used to analyse and quantify CD45^+^Ter119^+^ and CD45^-^Ter119^-^ cells by flow cytometry and representative FACS plots of these populations in control and infected mice **(D)** Quantification of absolute number of CD45^+^Ter119^+^ cells in control and infected mice. **(E)** Representative histogram plot demonstrating that CD45^+^Ter119^+^ cells express high levels of Sca-1 in response to *P. berghei*. **(F)** Quantification of the absolute number of CD45^+^Ter119^+^Sca-1^+^ cells in control and infected mice. **(G)** Quantification of absolute number of CD45^-^Ter119^-^ cells in control and infected mice. **(H)** Representative histogram plot demonstrating that CD45^-^Ter119^-^ cells express high levels of Sca-1 in response to *P. berghei*. **(I)** Quantification of the absolute number of CD45^-^Ter119^-^Sca-1^+^ cells in control and infected mice. **(C - I)** Data are representative of 3 independent infections. Error bars represent mean ± s.e.m.

To test the hypothesis obtained from the scRNAseq data, and to gain an indication of the strength of IFN signalling in the BM microenvironment, we measured the levels of IFN-γ in the serum and BM supernatant of control and infected mice. We focused on IFN-γ because this is the IFN type most widely associated with responses to *Plasmodium* infection in mice and humans (Meding *et al.*, 1990; John *et al.*, 2004; King & Lamb, 2015; Lelliott & Coban, 2016). Mice infected with *P. berghei* had elevated levels of IFN-γ in the serum and, in addition, we detected a striking increase of this cytokine in the BM (Figure 3B). Due to the fact that IFN levels were significantly high, and all HSPCs appeared to be affected by it, we hypothesized that not only HSPCs, but rather all BM cells, including stroma, would sense and respond to the elevated local and systemic IFN-γ. To test this, we used flow cytometry and assessed the extent of BM cells that may be responding to *P. berghei* infection, measuring Sca-1 upregulation at day 7 psi as a proxy of IFN-γ signalling (Ma, Ling & Dzierzak, 2001). Haematopoietic cells (CD45^+^ and/or Ter119^+^) were reduced in numbers, and, as expected, those that remained had substantially upregulated Sca-1 (Figure 3C-F). The overall number of CD45^-^ Ter119^-^ mixed haematopoietic and stroma cells (Boulais *et al.*, 2018) appeared to remain unchanged, however the majority of these cells upregulated Sca-1 (Figure 3C, G-I), as observed for HSPCs. These data suggested that IFN is affecting BM tissue globally and likely plays an important role in the response to infection, not only of haematopoietic cells but also stroma.

### *P. berghei* infection alters the integrity of vessels in the BM microenvironment

Having identified increased levels of IFN-γ within the BM and an IFN response from most BM cells, we set out to understand whether HSPC proliferation and loss of function may be linked to morphological and/or functional changes in the BM microenvironment. We started by studying endothelial cells (ECs), as they are known to support HSC maintenance and function (Hooper *et al.*, 2009; Kobayashi *et al.*, 2010; Butler *et al.*, 2010; Winkler *et al.*, 2012). We used IVM of mouse calvarium BM to identify changes in the distribution, morphology and function of BM ECs. To do this, we studied control and infected Flk1-GFP transgenic mice, in which CD45^-^ Ter119^-^ CD31^+^ phenotypic ECs express GFP and line BM blood vessels (Duarte *et al.*, 2018). Tilescans of BM calvarium revealed no dramatic changes in BM vessels (Figure 4A), and quantitative analysis of EC volume throughout the course of infection showed that the number of GFP^+^ voxels remained constant (Figure 4B). In agreement with this, flow cytometry analysis indicated that the number of ECs in long bones, including arteriolar Sca-1^+^ and sinusoidal Sca-1^-^ ECs, was unchanged by day 7 psi (Figure 4C-E).

**Figure 4:**
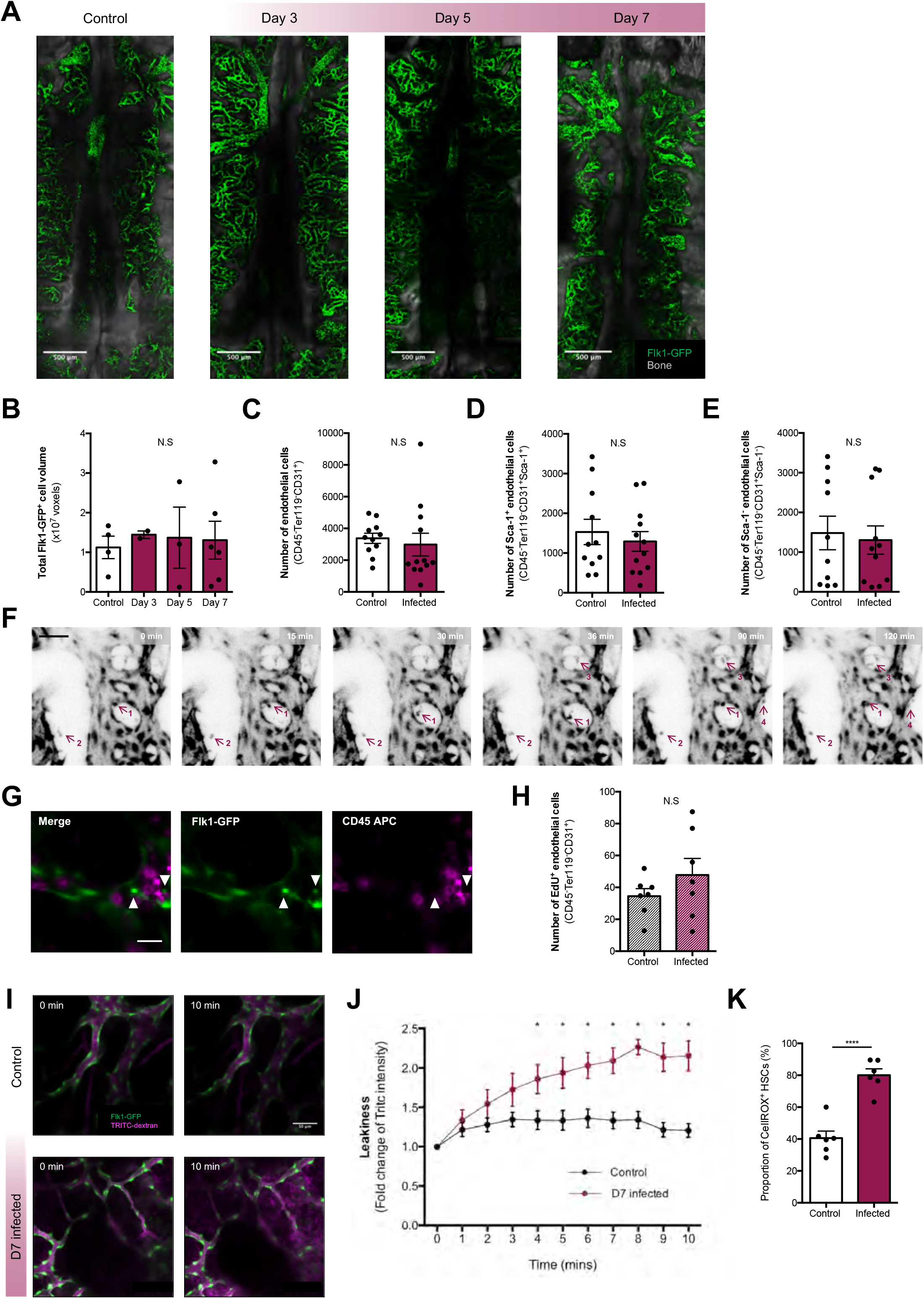
*P. berghei* infection affects vascular integrity. **(A)** Representative tilescan maximum projections (composite of individual tiles) of control and infected Flk1-GFP mice at day 3, 5 and 7 post-infection. Green, Flk1-GFP^+^ vessels; Gray, bone collagen SHG. **(B)** Automated segmentation and volume calculation (voxels) of control and infected Flk1-GFP^+^ ECs at day 3, 5 and 7 post-infection. **(C)** Absolute number of ECs (CD45^-^Ter119^-^CD31^+^) in control and infected mice at day 7 post-infection, analysed by flow cytometry. **(D)** Absolute number of arteriolar Sca1^+^ ECs (CD45^-^Ter119^-^CD31^+^Sca1^+^) in control and infected mice at day 7 post-infection, analysed by flow cytometry. **(E)** Absolute number of sinusoidal Sca1^-^ ECs (CD45^-^Ter119^-^CD31^+^Sca1^-^) in control and infected mice at day 7 post-infection, analysed by flow cytometry. **(F)** Selected frames from representative time-lapse data of an infected Flk1-GFP mouse at day 7 post-infection showing the detachment from the endothelium and migration of GFP^+^ particles indicated with purple arrow and numbered. Scale bar represents 150um. **(G)** Representative maximum projection of an area in an infected Flk1-GFP mouse at day 7 post-infection, injected with CD45 antibody. Green, Flk1-GFP^+^ vessels; Magenta, CD45 APC antibody. **(H)** Absolute number of EdU^+^ ECs in control and infected mice at day 7 post-infection. Data pooled from 2 independent infections. **(I)** Vascular leakiness was assessed by time lapse imaging of the calvarium for 10 minutes after injecting 3mg 65-85 kDa TRITC-dextran intravenously. Selected frames from representative time-lapse data of control and infected Flk1-GFP mice at day 7 post-infection showing the extravasation of vascular dye over time to allow quantification of vascular leakiness. **(H)** Quantification of leakiness in control and infected mice at day 7 post-infection quantified by measuring the fold change of TRITC intensity in 3 regions of interest per position. Data pooled from 4 independent infections. N = 4 control and 5 infected mice with 4 positions acquired per mouse. **(I)** Cellular ROS levels in HSCs analysed in control and infected mice. Data pooled from 2 independent infections. For **(B - E)** data pooled from 3 independent infections. For **(B-E, H, J and K)** error bars represent mean ± s.e.m.

Despite vessel morphology and EC numbers remaining unchanged, time-lapse IVM of randomly selected calvarium BM areas of Flk1-GFP mice revealed unusual dynamics over the course of a few hours. While the vasculature of control mice appeared essentially static (Supplementary Video 1), exclusively in infected animals, we detected GFP^+^ cells separating from those lining the vessel surface and migrating within the extravascular space with seemingly random trajectories (Figure 4F and Supplementary Video 2). To exclude that the migratory cells could be haematopoietic cells that expressed Flk1, we performed *in vivo* staining by injecting Flk1-GFP mice with APC-conjugated anti-CD45 antibody. The resulting data revealed that even though the size of GFP^+^ cells was similar to that of small CD45^+^ cells, they did not express CD45 themselves and therefore were not haematopoietic (Figure 4G). We next tested whether the observed GFP^+^ cells may be due to EC proliferation and flow cytometry analysis revealed no change in the number of EdU^+^ ECs by day 7 psi (Figure 4H).

As it has been shown that high levels of IFN-α leads to vascular leakiness (Prendergast *et al.*, 2016), and leakiness is a hallmark of stressed vasculature (Passaro *et al.*, 2017; Niz *et al.*, 2018), we next assessed whether vascular damage developed during the course of *P. berghei* infection. Time-lapse imaging of randomly selected calvarium BM areas of Flk1-GFP mice for a duration of 10 minutes following injection of low molecular weight TRITC-labelled dextran revealed that infected mice had highly permeable vessels, with dextran rapidly diffusing throughout the parenchyma (Figure 4I-J and Supplementary Video 3 and 4). This increased vascular leakiness was associated with a 2-fold rise in reactive oxygen species (ROS) in HSCs from infected animals (Figure 4K). These data suggested that increased vascular leakiness may affect HSC function.

### Dramatic loss of osteoblasts during *P. berghei* infection

Next, we focused on osteoblasts as they have long been associated with HSC function and maintenance of quiescence (Calvi *et al.*, 2003; Zhang *et al.*, 2003; Arai *et al.*, 2004; Lo Celso *et al.*, 2009), and we have recently demonstrated the importance of intact endosteal niches for healthy HSCs (Duarte *et al.*, 2018). Using IVM, we monitored osteoblast numbers and their distribution in control and infected Col2.3-GFP osteoblast reporter mice (Kalajzic *et al.*, 2002; Hawkins *et al.*, 2016). These experiments revealed a progressive and systemic loss of GFP^+^ cells over the course of infection to the extent that they were hardly detectable by day 7 psi (Figure 5A-B). To confirm that loss of GFP signal reflected loss of osteoblasts, we examined BM sections of femurs harvested from control and infected animals at day 7 psi, staining them with haematoxylin and eosin. While endosteum-lining cells were obvious in the sections of control mice (Figure 5C, black arrowheads), they were undetectable in those from infected animals (Figure 5C).

**Figure 5:**
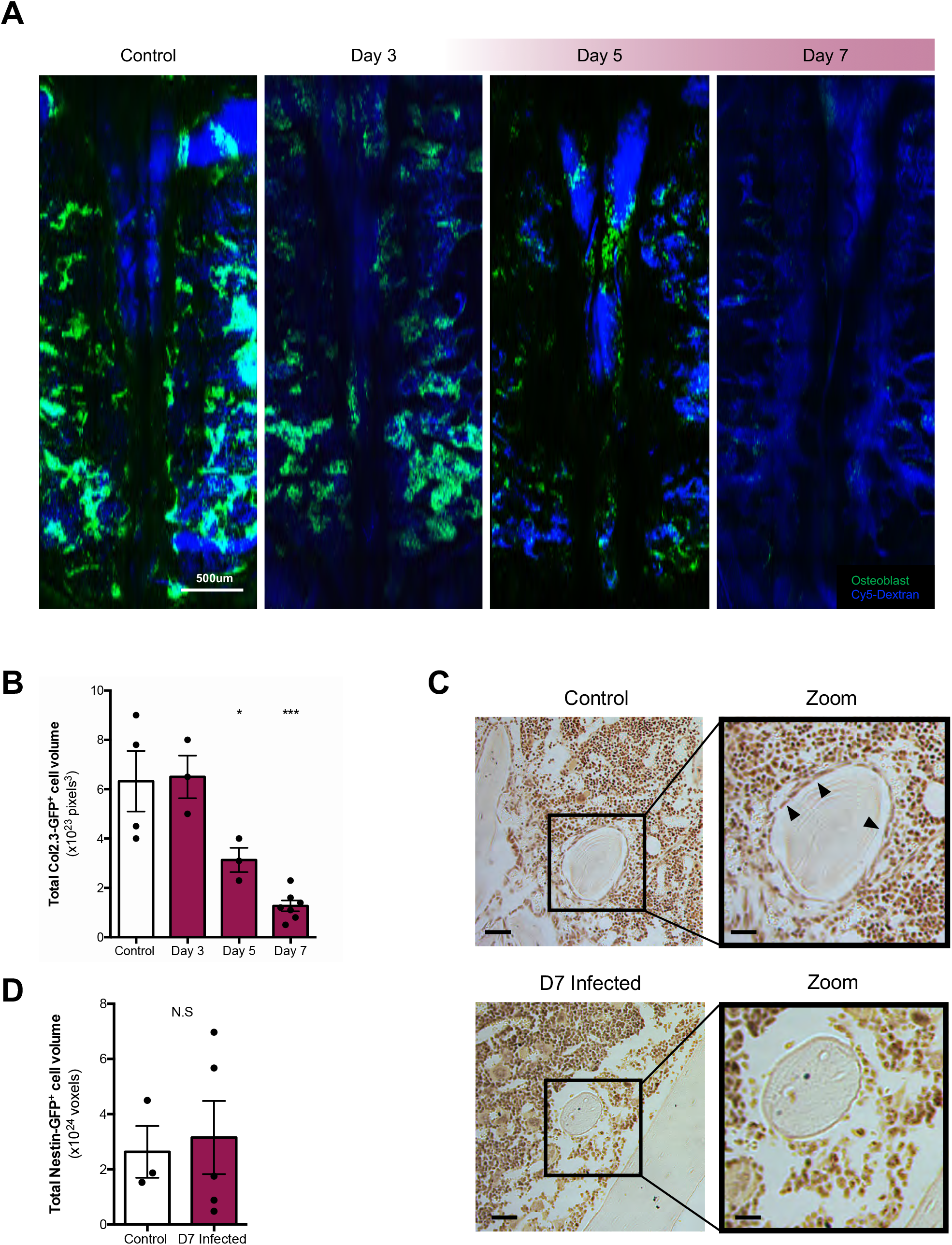
*P. berghei* infection causes a dramatic loss of osteoblasts. **(A)** Representative tilescan maximum projections (composite of individual tiles) of control and infected Col2.3-GFP mice at day 3, 5 and 7 post-infection. Blue, Cy5-dextran^+^ blood vessels; Green, Col2.3-GFP^+^ osteoblastic cells. **(B)** Automated segmentation and volume calculation (voxels) of control and infected Col2.3-GFP^+^ osteoblasts at day 3, 5 and 7 post-infection. Data were obtained from 5 independent infections. **(C)** Representative images of H & E stained sections of femurs from control and infected mice. Black arrowheads point at osteoblasts. Scale bars in original control and infected images represent 50um. Scale bars in control and infected zoom images represent 20um. **(D)** Automated segmentation and volume calculation (voxels) of control and infected Nestin-GFP^+^ MSCs at day 7 post-infection. Data were obtained from 2 independent infections. In **(C - D)** error bars represent mean ± s.e.m.

To understand whether primitive stroma cells may also be lost as a result of *P. berghei* infection, we studied Nestin-GFP mice in which GFP^+^ cells have been demonstrated to be stroma stem/progenitor cells upstream of osteoblasts (Méndez-Ferrer *et al.*, 2010). In these mice, however, the number of GFP^+^ cells remained unchanged by day 7 psi (Figure 5D), suggesting that malaria infection specifically targets mature osteoblasts.

### Maintaining osteoblasts inhibits HSC proliferation in response to *P. berghei* infection

Given the significant loss of osteoblasts observed within the BM of *P. berghei* infected mice, we hypothesized that this could be disrupting at least some HSC niches, and the mechanism triggering the observed increase in proliferation and loss of HSC function observed. We questioned whether manipulating the HSC niche by forcing the maintenance of osteoblasts through the course of infection could rescue HSCs. Parathyroid hormone (PTH) is a known activator of osteoblasts. Previous work has shown that intermittent treatment with PTH can enhance bone formation and increase osteoblast numbers by stimulating their proliferation and inhibiting apoptosis, and driving differentiation of osteoblast progenitors (Jilka *et al.*, 1999; Jilka, 2007; Jilka *et al.*, 2009; Kim *et al.*, 2012). We treated mice with PTH for two weeks prior to infection and for a further week as infection progressed before sacrifice and analysis (Figure 6A). While PTH treatment had no effect on parasitaemia (Supplementary Figure 2A), it did have an anabolic effect on osteoblasts in both control and infected mice. IVM of PBS (saline)- and PTH-treated, control and infected Col2.3-GFP mice at day 7 psi demonstrated that PTH treatment maintained a significant proportion of osteoblasts through the course of *P. berghei* infection (Figure 6B, C). The extent of the surviving osteoblast population was clear when we quantified osteoblast volume in all acquired tilescans and observed a significant, an approximately 3-fold increase in GFP^+^ voxels in infected PTH-treated mice vs. the control infected PBS treated group (Figure 6B-C). Of note, PTH treated, non-infected mice showed a significantly higher osteoblast volume than PBS treated mice (Figure 6C). To investigate the effects of PTH treatment on the dynamics of HSPC populations, we measured population size and proliferation rate of these cells in cohorts of PBS- and PTH-treated, control and infected mice. Consistent with Sca-1 upregulation occurring in both PTH and PBS infected mice (Supplementary Figure 2B), the proliferation rates and absolute numbers of LK cells and MPPs were similar in infected PBS- and PTH-treated mice (Figure 6D-F top and middle panels). Importantly, PTH treatment had an effect on the HSC response. Consistent with previous studies (Calvi *et al.*, 2003; Li *et al.*, 2012; Yao *et al.*, 2014), PTH treatment alone increased the number of LKS SLAM cells without increasing their proliferation rate. However, PTH-treated, infected mice did not show the typical increase in LKS SLAM cells proliferation at day 7 psi. Instead, these cells proliferated as little as those of control animals and significantly less than those of PBS-treated, infected mice (Figure 6D-F, bottom panels).

**Figure 6:**
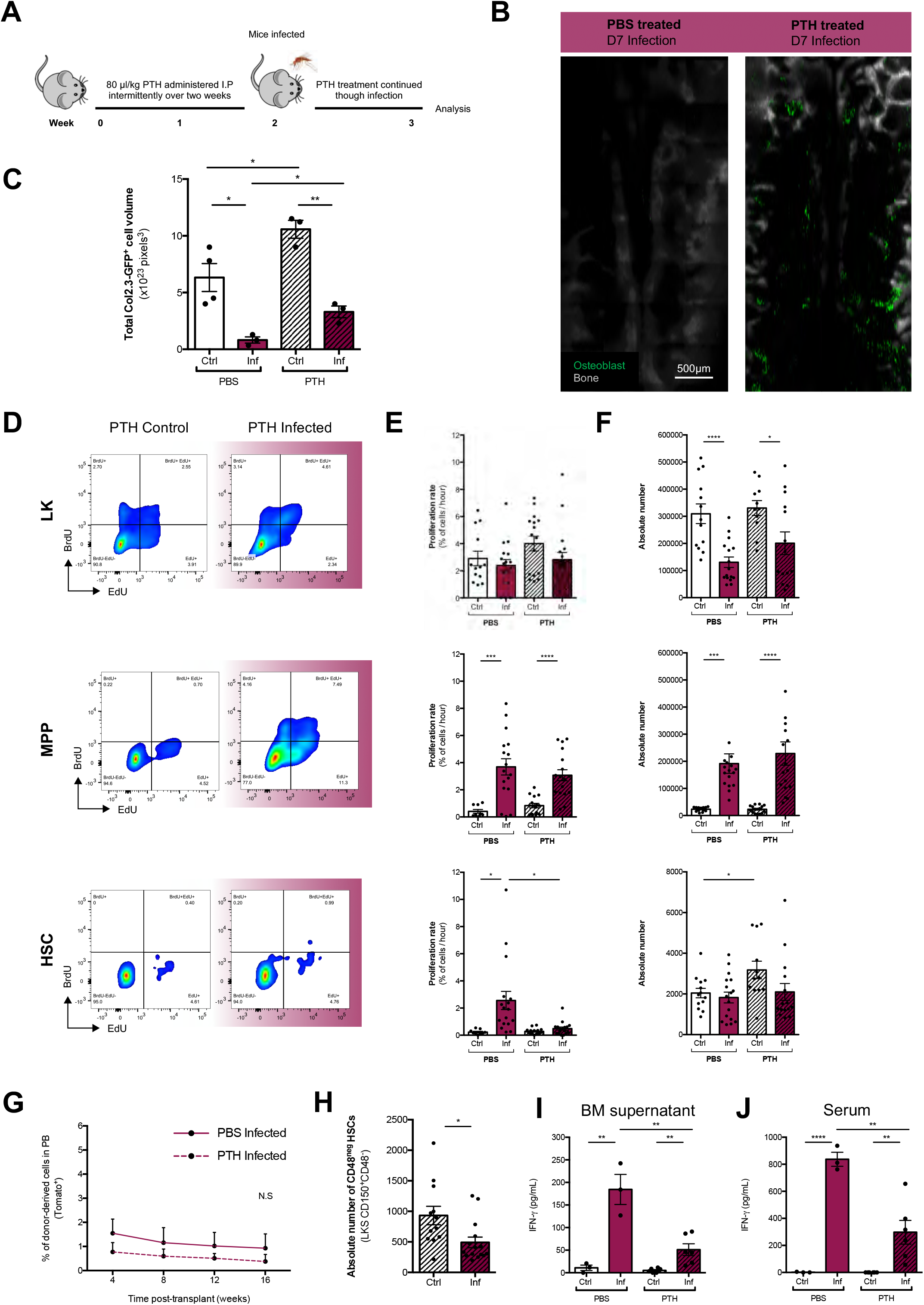
Parathyroid hormone treatment can reduce osteoblast loss, reduce overall IFN and proliferation of HSCs. **(A)** Scheme of treatment regimen used to reduce osteoblast loss through infection. **(B)** Representative tile scan maximum projections (composite of individual tiles) of PBS and PTH treated and infected Col2.3-GFP mice at 7 post-infection. Green, Col2.3-GFP^+^ osteoblastic cells. Gray, bone collagen SHG. **(C)** Automated segmentation and volume calculation (voxels) of PBS and PTH treated and infected Col2.3-GFP^+^ osteoblasts at 7 post-infection. Data were obtained from x independent infections. **(D)** Representative FACS plots showing the gating strategy for EdU^+^ and BrdU^+^ LK, MPP and HSC populations in PTH treated control and infected mice at day 7 post-infection. **(E)** Proliferation rates (proportion of cells cycling per hour) of LK, MPP and HSC populations analysed at day 7 post-infection in PBS and PTH treated control and infected mice. **(F)** Absolute numbers of LK, MPP and HSC populations analysed at day 7 post-infection in PBS and PTH treated control and infected mice. **(G)** Peripheral blood reconstitution assessed by flow cytometry post-primary transplant with HSCs from PBS and PTH treated infected donors at day 7 post-infection to lethally irradiated recipient mice. N = 6 mice per group. **(H)** Absolute number oh CD48-HSCs in PTH treated mice at day 7 post infection or control. **(I)** IFN-γ levels in BM supernatant of PBS and PTH treated control and infected mice measured by ELISA. N = 3 mice per PBS group and N = 6 mice per PTH group. **(J)** IFN-γ levels in serum of PBS and PTH treated control and infected mice measured by ELISA. N = 3 mice per PBS group and N = 6 mice per PTH group. For **(C, E – J),** Error bars represent mean ± s.e.m.

To test whether PTH treatment could also rescue HSC function, we performed transplantation assays in which we compared the reconstitution ability of primitive HSPCs purified from PBS- and PTH-treated, infected donors at day 7 psi. We observed no difference between the two groups and the overall engraftment was very low (Figure 6G), indicating that PTH treatment did not rescue HSC function. Consistent with this, the number of CD48^-^ HSCs in PTH treated, infected mice was lower than in non-infected ones (Figure 6H). To understand why HSC function was not rescued by PTH treatment, we analysed the levels of IFN-γ in the BM supernatant and serum of PBS- and PTH treated control and infected mice at day 7 psi. Interestingly, the concentration of IFN-γ in both the BM and serum of PTH-treated and infected mice was significantly lower than that of PBS-treated and infected mice. However, IFN-γ was, however, still augmented in PTH treated infected animals compared to PTH-treated controls, albeit to a lesser extent than in infected PBS-treated mice (Figure 6I-J). Together, these data suggest that although PTH treatment leads to the retention of osteoblasts, inhibition of HSC proliferation and reduction of both systemic and localised IFN-γ levels, the remaining IFN-γ is still sufficient to affect the long-term reconstituting capacity of HSCs exposed to *P. berghei*, through direct or indirect action on HSCs.

## DISCUSSION

HSCs maintain blood production thanks to complex interactions with multiple components of the BM microenvironment, however, their function can be affected by severe infections. Here we used an experimental model of *Plasmodium* infection to gain a quantitative understanding of the impact of malaria disease on HSPC proliferation and function and to investigate the role of the BM microenvironment in mediating the phenotypes observed. Sporozoite-mediated *P. berghei* infection was chosen as our model as it is a natural rodent infection that includes the skin and livers stages and reproduces several features of human malaria. Malaria is a widespread and severe infection and its effects on HSCs are still poorly understood beyond reports of increased morbidity and infection susceptibility in survivors (Orf & Cunnington, 2015; White, 2018).

We measured the proliferation rate of HSPC populations during the course of infection using a dual-pulse labelling method that we recently adapted to the haematopoietic system, based on sequential injections of EdU and BrdU, followed by flow cytometry analysis (Akinduro *et al.*, 2018). Consistent with our previous analyses of early haematopoiesis during *P. berghei* infection (Vainieri *et al.*, 2016) and with studies of other experimental models of severe infection (Baldridge *et al.*, 2010a; Matatall *et al.*, 2016; Vainieri *et al.*, 2016), we detected increased proliferation of primitive HSPCs and increased proliferation and compartment size of MPPs. While previous observations were based on the incorporation of a single nucleotide and, therefore, only provided qualitative information, the data generated here could be directly entered into simple mathematical models that demonstrated the extent of increased turnover of both the HSC and MPP compartments. In particular, we could conclude that the increase in size of the MPP compartment in infected mice could be driven solely by HSC and MPP proliferation, and that by day 5 psi, a significant proportion of MPP progeny must exit the compartment.

Emergency myelopoiesis has been described to be induced by multiple infections (MacNamara *et al.*, 2011; Boettcher *et al.*, 2012; Schürch, Riether & Ochsenbein, 2014). In line with this, our scRNAseq analysis highlighted skewing of early haematopoiesis towards the generation of myeloid immune cells at the expense of erythroid and megakaryocyte differentiation, together with a loss of the most primitive HSPC transcriptional signature. Modelling and scRNAseq data were consistent with the loss of HSC function that we observed in transplantation assays. While our data suggest that the overall HSC compartment turned over more than once over the course of infection, the question remains whether this is driven by the fast proliferation of a specific subset of HSCs or by slower proliferation of the entire compartment. In other words, are all HSCs equally affected by infection, or is there a subset that responds more readily, allowing for another subset to remain relatively untouched? Given that: 1) some cells remain that express the most primitive signature; 2) phenotypically defined HSC subsets enriched for the highest functionality (E-SLAM and CD48^neg^ HSCs) are reduced but not lost; and 3) primary HSC transplants show a significant loss of function, but primitive HSPCs purified from primary recipients of control and infected HSCs, respectively, performed similarly together suggest that a small fraction of HSCs may survive infection relatively unscathed. These are likely the CD48^-^/EPCR^+^ subsets of the LKS SLAM population.

Gene expression profiling of stem and progenitor cell populations highlighted the central role of interferon in driving the observed phenotypes. IFN-γ is highly expressed in response to multiple strains of human and murine *Plasmodium* and it has an important role in controlling the strength of infection (Meding *et al.*, 1990; John *et al.*, 2004; Lelliott & Coban, 2016). It has, however, also been reported to mediate and exacerbate the disease itself, causing damage to the host (King & Lamb, 2015). We uncovered that IFN-γ damages not only early haematopoiesis, but also BM stroma, which are known components of the HSC niche. This is likely due to its accumulation within the tissue. BM vasculature damage was recently reported in response to IFN-γ (Prendergast *et al.*, 2016) and in mice burdened with malaria, where it was proposed to mediate parasite accumulation in the tissue (Niz *et al.*, 2018). Our data on vascular leakiness and HSC ROS levels are consistent with previous observations that steady-state HSCs residing in the vicinity of more permeable vessels contain higher levels of ROS and are less functional (Itkin *et al.*, 2016), suggesting that increased vascular permeability may be one of the mechanisms leading to loss of HSC function during *P. berghei* infection. Due to the fact that parasite accumulation in the BM has been reported in both mice and humans (Joice *et al.*, 2014), and vascular damage is likely to be similar in both species, it is possible that the same mechanism affects HSC function in both as well.

It was recently reported that both *Plasmodium* infection and sepsis affect bone biology (Terashima *et al.*, 2016; Lee *et al.*, 2017) and increasing attention is being given to a potential role of the BM microenvironment in maintaining a reservoir of *Plasmodium* parasites in human malaria (Aguilar *et al.*, 2014; Joice *et al.*, 2014; Niz *et al.*, 2018). In particular, sepsis-induced loss of osteoblasts has been linked to the loss of lymphoid progenitors and haematopoiesis being skewed towards myeloid lineages. This mechanism is likely to take place in response to *Plasmodium* infection too. In addition, we recently showed that loss of osteoblasts is linked to the localised loss of HSCs driven by leukaemia (Duarte *et al.*, 2018). In contrast, in P. *berghei* infection, loss of osteoblasts is specifically linked to primitive HSPC proliferation and loss of function, likely through loss of CD48^-^ and EPCR^+^ HSC subsets. Osteoblast rescue through PTH treatment eliminated HSC proliferation but did not preserve HSC function. This is an important finding as HSC proliferation and loss of function are usually directly linked (MacNamara *et al.*, 2011; Matatall *et al.*, 2016), but they were uncoupled here. This loss of function is most likely linked to the persistence, albeit at reduced levels, of IFN-γ both systemically and within the BM microenvironment. The reduced levels of IFN-γ are potentially due to the immunomodulatory properties of PTH. It has been described that T cells contribute to the anabolic processes triggered by this hormone (Terauchi *et al.*, 2009; Li *et al.*, 2012), and it is likely that PTH affects their response to *Plasmodium* infection. Importantly, haematopoietic damage in response to IFN has traditionally been associated with HSC proliferation, but our study shows that it is possible to uncouple HSC proliferation and loss of function, suggesting that IFN exposure leads to the loss of HSC function through additional mechanisms. Increased vascular leakiness and ROS accumulation may be two of them. While the data presented here shows that manipulating osteoblasts could be a way of reducing the damage caused by malaria, future work will focus on finding alternative, complementary approaches to reduce the damage induced by IFN. These will be important to curb long-term stem cell damage in survivors of malaria and other severe infections.

## METHODS

### Mice

All animal work was in accordance with the animal ethics committee (AWERB) at Imperial College London and UK Home Office regulations (ASPA, 1986). Flk1-GFO mice were a gift from Alexander Medvinsky (University of Edinburgh, (Xu *et al.*, 2010). Tuck CD1 and C57BL/6 mice were obtained from Envigo or Charles River. Col2.3-GFP (Hawkins *et al.*, 2016), mT/mG (Muzumdar *et al.*, 2007) and CD45.1 (BJ/SJL) mice were bred and housed at Imperial College London, according to Institutional guidelines. Male and female mice greater than 6 weeks old were used in all experiments.

For PTH treatment of mice for the maintenance of osteoblasts, Rat PTH (1-34) (Bachem) (80µg/mg/kg body weight) or PBS (vehicle control) was injected intraperitoneally (i.p) five times a week for two weeks prior to the infection and for a further week through the infection until mice were sacrificed for analysis. For these experiments, 6-week-old female mice were used.

### *Plasmodium berghei* experimental model

Parasite maintenance, generation of infected mosquitos and infection of mice were carried out as previously described (Ramakrishnan *et al.*, 2013; Vainieri *et al.*, 2016). Briefly, *P. berghei* ANKA 2.34 was maintained in 4-10-week-old female Tuck CD1 mice by serial blood passage and according to Home Office approved protocols. Hyper reticulocytosis was induced 2-3 days before infection by treating mice with 6mg/ml phenylhydrazine chloride. Stock mice were infected by intraperitoneal injection of blood containing parasites and parasitemia was monitored on Giemsa-stained tail blood smears and expressed as a percentage of more than 500 RBCs counted per slide. For each individual experiment, a group of five female CD1 phenylhydrazine chloride treated mice were infected with *P. berghei*, injected i.p, followed by feeding to mosquitos. Infected mice are subsequently anaesthetized and exposed to cages containing 500 starved female *Anopheles stephesi* mosquitos. Fed mosquitos were maintained until 21 days post infection, at which point salivary gland sporozoites are at their peak. To infect mice with *P. berghei* for experiments, each mouse was exposed to 5 infected mosquitos and control mice receive 5 mosquito bites from naïve, non-infected mosquitos in parallel. Blood parasitemia and symptoms of malaria were monitored and mice sacrificed for analysis at various timepoints of interest and before the onset of cerebral malaria.

### Transplantation assays

Tibias, femurs, ileac bones, vertebrae and sternum were harvested from PBS or PTH treated control and infected mT/mG or CD45.1 donor mice, The bones were crushed, filtered through a 40um strainer and the red blood cells lysed. Whole BM was labelled with a cocktail of biotinylated lineage antibodies (CD3, CD4, CD8, Ter119, B220, Ly6G, CD11b) and streptavidin magnetic Microbeads (Miltenyi Biotech) to perform a lineage depletion using the MACS Column Technology (Miltenyi Biotech). The lineage depleted sample was then stained and sorted for HSCs (defined as Lineage^-^ c-Kit^+^ Sca-1^+^ CD48^-^ CD150^+^). 200 HSCs were transplanted into lethally irradiated WT recipients (two doses of 5.5Gy, at least three hours apart) alongside 200,000 competitor WT BM cells. Baytril antibiotic was administered to recipient mice for 6 weeks post-transplant. Multilineage engraftment of the recipients was monitored every 4 weeks post-transplant up to week 20 by FACS analysis of the peripheral blood. For secondary transplants, whole BM was harvested from primary recipients of control or infected HSCs, pooled and re-sorted for HSCs and transplanted into s second cohort of lethally irradiated WT recipients, with each mouse again receiving 200 HSCs and 200,000 competitor WT BM cells. Multilineage engraftment was monitored as with the primary recipients.

### Flow Cytometry

For the analysis of haematopoietic stem and progenitor cell compartments, bones were crushed in PBS with 2% fetal bovine serum (2%FBS) and the cells were filtered through a 40um strainer, depleted of red blood cells and stained with relevant antibodies. For stroma analysis, tibias and femurs were crushed, digested with collagenase I (Worthington, UK) at 37C, for 20 minutes with 110rom agitations. The cells obtained were filtered through a 70um strainer and stained with relevant antibodies. For the analysis of peripheral blood, approximately 10ul of blood was obtained from mice via the tail vein and mixed with EDTA to prevent clotting. Red blood cells were lysed, the cells washed with 2%FBS and subsequently stained with relevant antibodies. The following fluorochrome-conjugated or biotinylated primary antibodies specific to mouse were used: CD3, Ter119, B220, CD11b, cKit, Sca-1, CD150, CD48, CD31, all from Biolegend, and EPCR from StemCell Technologies. For secondary staining, streptavidin Pacific Orange (Invitrogen) was used. Live and dead cells were distinguished using DAPI (Invitrogen) or, where necessary, a fixable viability dye (LifeTechnologies). BrdU and EdU were detected and analysed as previously described (Akinduro *et al.*, 2018) using a combination of the BrdU kit (Becton Dickinson) and the Click-iT EdU Kit (Life Technologies). For the analysis of cellular ROS, CellROX Deep Red Reagent was used, following manufacturer’s instructions (ThermoFisher Scientific). Calibrite beads (BD Biosciences) were used to determine the absolute cell numbers in the populations of interest, as described previously (Hawkins *et al.*, 2007). Cells were analysed with a LSR-Fortessa (BD Biosciences) and data were analysed with FlowJo (Tree Star).

### Mathematical modelling

Simple mathematical models for the progression of the HSC and MPP cell populations in control and infected mice were developed in order to extrapolate from empirical data. We use the following notation for t in {1,2,3,5,7} days: M_ctrl_ denotes the total number of control mice measured; N^HSC^_ctrl,i_(t) denotes the number of HSCs measured in control mouse i at time t; N^MPP^_ctrl,i_(t) denotes the number of MPPs measured in control mouse i at time t; S^HSC^_ctrl,i_(t) denotes the number of HSCs entering S-phase per hour measured in control mouse i at time t; and S^MPP^_ctrl,i_(t) denotes the number of MPPs entering S phase per hour measured in control mouse i at time t. We let M_inf_, N^HSC^_inf,i_(t), N^MPP^_inf,i_(t), S^HSC^_inf,i_(t), and S^MPP^_inf,i_(t) denote the equivalent measurements in infected mice. In addition we define M_inf_(3) to be the number of infected mice up to day 3 and M_inf_(5,7) to be the total number of infected mice measured in days 5 and 7.

As the number of HSCs in both the control and infected mice are consistent over all five days of measurement, in both the control and infected model their number is set to a constant, the average value, corresponding to the maximum likelihood estimate,

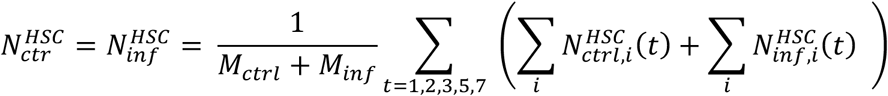

for all t. As the numbers of MPPs in the control mice are consistent with those up to day three in the infected mice, the control model is

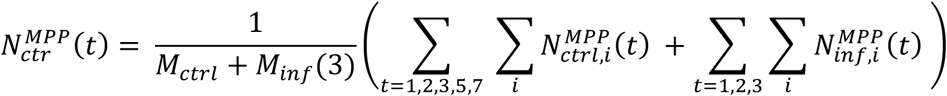

for all t. In the infected model, the number of MPPs coincides with the number in the control model up until day 3, 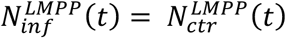 for t up to day 3. How that increases thereafter depends on the increased proliferation and the differentiation assumption as described below.

The number of HSCs entering S-phase per hour is assumed consistent between the control mice and the infected mice until day 3 so that

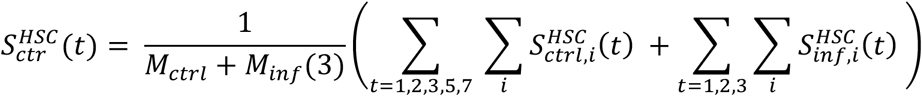

and 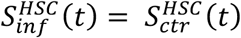 for t up to day 3. In the infected model, the number of HSCs entering S phase per hour is assumed constant for t greater than 5 days at the average value

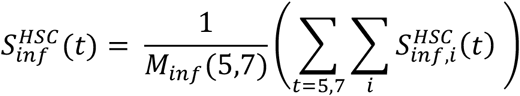

Between days 3 and 5, in the infected model, we assume a linear increase in the per-hour rate of entry into S-phase of HSCs from the control to infected proliferation:

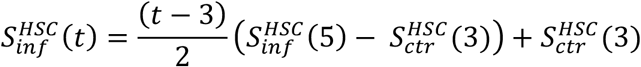

for t in (5,7). Analogous assumptions are made for MPP proliferation with the infected and control model concurring until day 3, whereupon the infected model experiences a linear increase in number of cells entering S-phase per hour to the average of the 5 and 7 empirical values at day 5, then becoming constant.

The maroon line in Figure 1G is constructed assuming that the additional output caused by the increased number of HSCs entering S phase per hour beyond that in the control mouse become MPPs,

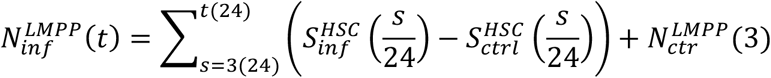

The dashed maroon line in Figure 1G is constructed assuming that the additional output caused by the increased entry into S phase of both HSCs and MPPs beyond their control equivalents all become MPPs:

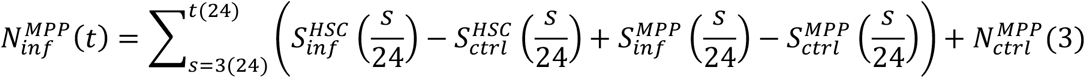

The model lines in Figure 1H are constructed by considering the number of cells that have entered S phase over time in each of the models as a proportion of the control compartment size. Namely, for the control HSC model we have

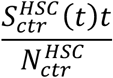

while for the infected HSC model the proportion is

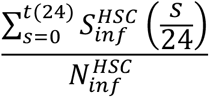

Analogous expressions are used for MPP model.

### Single cell RNA-seq and analysis

For 10x Chromium™ (10x Genomics, Pleasanton, CA) experiments, Lineage-c-Kit+ cells were sorted and processed according to the manufacturer’s protocol.

*Data processing and Quality Control*: sample demultiplexing barcodes processing and gene counting was performed using the count commands from the Cell Ranger v1.3 pipeline. Each sample was filtered for potential doublets by simulating doublets from pairs of scRNAseq profiles and assigning scores based on a k-nearest neighbour classifier on PCA transformed data. The top 4.5% of cells with the highest doublet scores from each sample were removed from further analysis. Cells with >10% of unique molecular identifier (UMI) counts mapping to mitochondrial genes, expressing fewer than 500 genes, or with the total number of UMI counts further than 3 standard deviations from the mean were excluded.

*Identification of variable genes:* variable genes were identified for all samples following the method of Macosko *et al*. with expression = 0.001 and dispersion = 0.05 used as minimum cut-offs. Cells were normalised to the same total count for each cell and log transformed (x -> log(x+1)). Each gene was scaled so it was zerocentered. Typically, 5000 variable genes were carried forward for analysis. All further analysis was performed using the Scanpy Python module. In total there are 14193 control and 13905 infected cells.

*Clustering and mapping:* The control cells were clustered using Louvain clustering (Scanpy igraph method). To assign infected cells to clusters, the infected samples were projected into the PCA space of the control data and the nearest neighbours calculated between the control and infected samples based on the Euclidean distance in the top 50 components. Infected cells were assigned to the same cluster that the majority of their 15 nearest control neighbours belonged to.

Force graphs for the control and infected data sets were created by constructing a k = 7 nearest neighbour graphs on the top 50 principal components of the variable genes for each dataset. Edge lists were exported into Gephi 0.9.1 and graph coordinates calculated using the ForceAtlas 2 layout.

The bar graph showing the abundance changes of certain cell types between control and infected were created by calculating the percentage abundances of each cell cluster and calculating the log2-fold change of the infected cells compared to control. The infected cells were split into two samples, each corresponding to one mouse.

#### Binary clustering

The binary clustering was created by choosing the control clusters that showed clear expression of genes known to associate with a HSC state, including *Procr* and *Sca-1*. Clusters 0 and 13 were chosen to contain the most immature cells and were therefore distinguished from the others in blue. The infected samples were split in a similar fashion, based on the cells that mapped to clusters 0 and 13 in the control data.

#### Determining initial IFN genes and correlating IFN genes

Differential expression was performed between the control and infected cells for each cluster. This was done in the R package edgeR, using a ratio likelihood test. Genes that were at least 4-fold upregulated in the infected compared to WT in all clusters were chosen as an initial set of driving genes involved in the infection. This list contained 20 genes. The entire set of variable genes (calculated using the combined control and infected samples) was analysed to find all the genes whose expression correlated highly with any of those in the initial set of 20. Genes that had a spearman correlation > 0.3 with any of these genes were added to the list of driving genes. This procedure added 89 genes for a total of 109 driving genes.

### Enzyme-linked immunosorbent assay (ELISA)

To obtain BM supernatants, tibias and femurs were harvested from control and infected mice. The metaphysis and diaphysis of long bones were separated with scissors and 80ul of PBS was flushed through each diaphysis, collected and re-flushed. Cells were pelleted by centrifugation at 400 g for 5 minutes. The supernatant was collected and any remaining cells excluded by centrifugation at 500 g for 5 minutes. Serum was prepared by collecting blood by cardiac puncture after terminal anaesthesia with pentobarbital. Blood was incubated at 4C for at least 3 hours to clot and subsequently centrifuged at 12,000 g for 10 minutes at 4C. The serum supernatant was transferred to a new Eppendorf tube. BM supernatants and serum were stored at −20C until required for ELISA. IFN-g ELISAs (Biolegend) were performed according to the manufacturer’s instructions and by diluting the BM supernatant 1:4 and serum samples 1:2.

### Intravital microscopy

Intravital microscopy was performed using a Zeiss LSM 780 upright confocal microscope equipped with Argon (458, 488 and 514nm), a diode-pumped solid state 561 nm laser and a Helium-Neon 633 nm, a tunable infrared multiphoton laser, 4 non-descanned detectors (NDD) and a n internal spectral detector array. Live imaging of the calvarium BM was carried out as described in (Hawkins *et al.*, 2016; Duarte *et al.*, 2018). Blood vessels were labelled with 80µl of 8mg/ml Cy5-Dextran (nanocs, MA). Second harmonic signal was excited at 860-880nm and detected with external detectors. CFP signal was excited at 870nm or 458nm and detected using external or internal detectors; GFP signal: excitation at 880nm or 488nm, external or internal detectors; YFP signal: excitation at 488nm or 514nm, internal detectors. mTomato/DsRed and Cy5 signals were respectively excited at 561nm and 633nm and detected using internal detectors. Assessment of vasculature leakiness with infection was carried out by adapting the protocols from (Itkin *et al.*, 2016; Passaro *et al.*, 2017).

### Bone marrow histology

Femurs were harvested from infected and control animals at day 7 psi, fixed in 4% PFA at 4°C overnight and decalcified in 10% EDTA for 3 days. The bones were subsequently embedded in paraffin, sectioned and processed for haematoxylin and eosin staining at the NHLI histopathology facility at Imperial College.

### Image processing and quantification

Zen black (Zeiss, Germany) software was used to stitch three-dimensional BM tilescans (tilescans represent individual tiles stitched together to form a composite). ImageJ was used to visualise, register and process raw data. Fiji was used to manually crop out autofluorescent signal out of the tissue. Automated cell segmentation and volume measurements were performed in Definiens (Definiens Developer 64, Germany). For analysis of TRITC-dextran extravasation in time lapse movies from control and infected mice, pixel intensity within 4 equally sized and randomly placed regions of interest was calculated per time frame. The average of these values was subsequently calculated and the fold change increase in intensity in control and infected samples plotted in the graph.

### Quantification and statistical analysis

Raw data was visualised and processed using Microsoft Excel, MATLAB and GraphPad Prism (GraphPad Software Inc.). Group means were compared using the unpaired Student’s t test. For multiple comparisons, one-way ANOVA with post hoc Tukey test or Bonferroni correction was used. For all data, differences were considered significant whenever p<0.05. *p<0.05; **p<0.01; ***p<0.001; ****p<0.0001. Specific statistical details including the number of animas used can be found in the figure legends.

## Supporting information

## ACKNOWLEDGMENTS

This work was funded by ERC (ERC STG 337066 to CLC), BBSRC (BB/L023776/1 to CLC), the Wellcome Trust (PhD studentship to MLRH). AMB was funded by the MRC (New Investigator Research Grant MR/N00227X/1. Work in the Gottgens group is funded by the Wellcome Trust, Bloodwise, CRUK and core funding by the wellcome and MRC to the Wellcome & MRC Cambridge Stem Cell Institute. SW is the recipient of an MRC studentship. CLC and KD were supported in part by the Royal Irish Academy – Royal Society International Exchange Cost Share Program (IEC\R1\180061).

We are particularly grateful to Mark Tunnicliffe for passaging parasites and preparing mosquitos for the experiments performed and to Dr Fiona Angrisano and Kasia Sala for technical support with infections. We also thank the Imperial College DoLS Flow Cytometry facility and Central Biomedical Services for logistical support as well as Jean Langhorne, Helen Fletcher and Aubrey Cunnington for constructive discussions.

## Supplementary Figure and Video legends

**Supplementary Figure 1.**
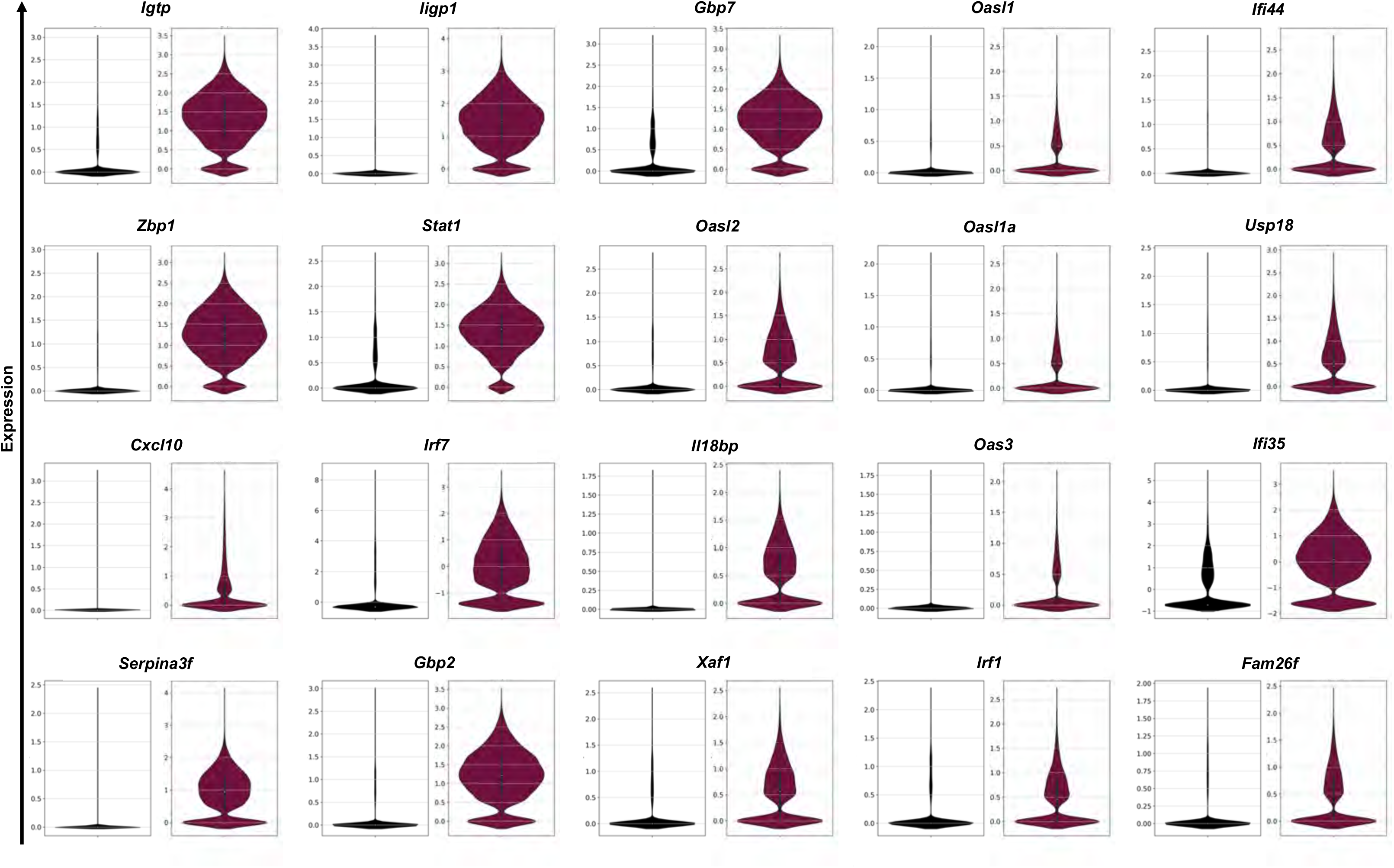
Violin plots showing the expression of the 20 driving genes determined via scRNAseq analysis in control (black, left panels) and infected (marron, right panels) mice.

**Supplementary Figure 2.**
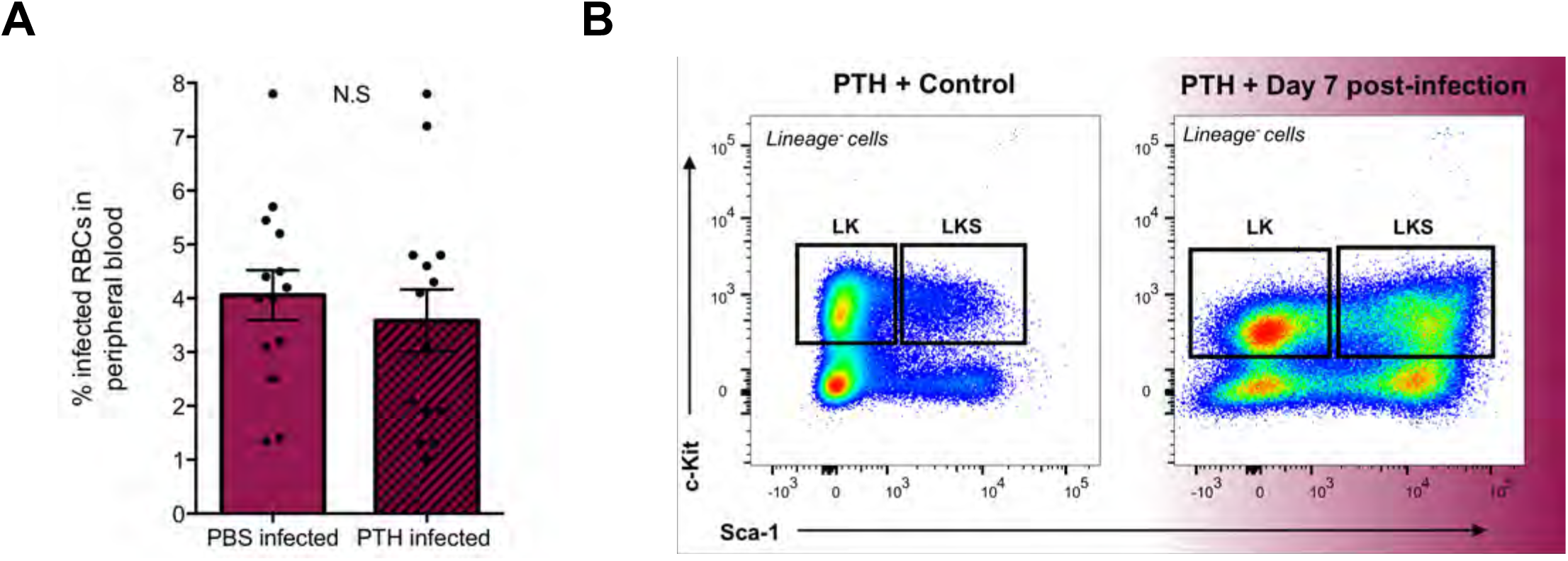
**(A)** Parasitemia in peripheral blood calculated from Giemsa-stained blood films taken from PBS- and PTH-treated mice at day 7 psi and quantified. Each dot represents one mouse. **(B)** Representative FACS plots showing the changing pattern of Sca-1 and c-Kit expression in Lineage^-^ BM haematopoietic cells in PTH treated control and infected mice at day 7 psi. Boxes indicate the gating strategy used for LKS and LK populations.

**Supplementary Table 1.** The 109 driving genes identified by scRNA-Seq analysis.

| Driving Genes |  |  |  |
| --- | --- | --- | --- |
| Igtp | Zbp1 | Cxcl10 | Serpina3f |
| Iigp1 | Stat1 | Irf7 | Gbp2 |
| Gbp7 | Oasl2 | Il18bp | Xaf1 |
| Oasl1 | Oas1a | Oas3 | Irf1 |
| Ifi44 | Usp18 | Ifi35 | Fam26f |
| 1700016F12Rik | AW112010 | Apobec3 | Apol9b |
| B2m | Bst2 | Calr | Capza2 |
| Cd74 | Cebpb | Csf2rb | Ctsz |
| Cxcl10 | Dhx58 | Dtx3l | Epsti1 |
| F830016B08Rik | Fam26f | Ffar2 | Gbp2 |
| Gbp3 | Gbp4 | Gbp5 | Gbp7 |
| Gm12185 | Gm4841 | Gm4951 | H2-Aa |
| H2-Ab1 | H2-D1 | H2-DMa | H2-DMb1 |
| H2-Eb1 | H2-K1 | H2-Q6 | H2-Q7 |
| H2-T22 | H2-T23 | Hsp90aa1 | Hspa5 |
| Hspe1 | Ifi203 | Ifi204 | Ifi27l2a |
| Ifi35 | Ifi44 | Ifi47 | Ifit1 |
| Ifit3 | Ifitm1 | Ifitm3 | Irgm1 |
| Isg15 | Isg20 | Lap3 | Lgals3bp |
| Lgals9 | Ly6a | Ly6c2 | Ly6e |
| Manf | Mif | Mnda | Mndal |
| Ms4a4c | Ms4a6b | Ms4a6c | Nmi |
| Pacsin1 | Parp12 | Parp14 | Parp9 |
| Pla2g16 | Plac8 | Ppa1 | Psmb10 |
| Psmb8 | Psmb9 | Psme1 | Pycard |
| Rnf213 | Rsad2 | Rtp4 | Samhd1 |
| Sdf2l1 | Serpina3g | Shisa5 | Slfn2 |
| Socs1 | Tap1 | Tapbp | Trafd1 |
| Trim12c | Trim30a | Ube2l6 | Usp18 |

**Supplementary Video 1**

Representative maximum projection of 3D time-lapse data (shown at 10 frames per second) of an area from a control Flk1-GFP mouse collected every 90s for 120min. Representative of 4 control mice.

**Supplementary Video 2**

Representative maximum projection of 3D time-lapse data (shown at 10 frames per second) of an area from an infected Flk1-GFP mouse at day 7 psi collected every 90s for 120min. Representative of 7 infected mice.

**Supplementary Video 3**

Representative maximum projection of 3D time-lapse data (shown at 10 frames per second) of an area from a control Flk1-GFP mouse injected with low molecular weight TRITC-dextran collected every minute for 10 minutes to assess vascular leakiness. Green: Flk1-GFP ECs; magenta: TRITC dextran. Representative of 4 control mice.

**Supplementary Video 4**

Representative maximum projection of 3D time-lapse data (shown at 10 frames per second) of an area from an infected Flk1-GFP mouse at day 7 psi injected with low molecular weight TRITC-dextran collected every minute for 10 minutes to assess vascular leakiness. Green: Flk1-GFP ECs; magenta: TRITC dextran. Representative of 5 infected mice.

